# Benchmarking the Impact of Anatomical Segmentation on In Vivo Magnetic Resonance Spectroscopy

**DOI:** 10.1101/2025.07.15.664931

**Authors:** Jessica Archibald, Kay Chioma Igwe, Antonia Kaiser, Karl Landheer, Jaimie Lee, John L.K. Kramer, Niklaus Zölch, Aaron Gudmundson, Helge J. Zöllner, Candace C. Fleischer, Georg Oeltzschner, Jamie Near, Mark Mikkelsen

**Affiliations:** Department of Radiology, Weill Cornell Medicine, New York, New York, USA; Department of Biomedical Engineering, Columbia University Fu Foundation School of Engineering and Applied Science, New York, NY, USA; CIBM Center for Biomedical Imaging, Ecole Polytechnique Fédérale de Lausanne, Lausanne, Switzerland; Regeneron Genetics Center, Tarrytown, NY, USA; Department of Anesthesiology, Pharmacology and Therapeutics, Faculty of Medicine, University of British Columbia, Vancouver, BC, Canada; Institute of Forensic Medicine Universität Zürich, Zürich, Switzerland; Russell H. Morgan Department of Radiology and Radiological Science, The Johns Hopkins University School of Medicine, Baltimore, MD, USA; Department of Radiology and Imaging Sciences, Emory University School of Medicine, Atlanta, GA, USA; Department of Biomedical Engineering, Georgia Institute of Technology and Emory University, Atlanta, GA, USA; Sunnybrook Research Institute and University of Toronto, Toronto, Canada

**Keywords:** ANTS, brain metabolites, FSL, MR spectroscopy, SPM, quantification, tissue segmentation

## Abstract

**Purpose:** Estimation of metabolite concentrations in brain magnetic resonance spectroscopy (MRS) requires correction for differences in tissue water content, relaxation properties, and the proportions of gray matter (GM), white matter (WM), and cerebrospinal fluid (CSF). Accurate knowledge of the relative proportions of these tissue classes within the volume of interest is therefore essential for reliable quantification. Commonly used brain segmentation tools differ in their algorithms, priors, and implementation, potentially introducing variability in MRS-derived concentration estimates. This study investigates the impact of segmentation software on estimated absolute concentrations.

**Methods:** Three segmentation software tools, ANTs, FSL, and SPM, were evaluated. Segmentations were applied to an in vivo test-retest MR dataset to assess (1) differences in estimated tissue fractions, and (2) how these differences propagate into tissue-corrected metabolite concentrations. As an additional validity check and biological benchmark of segmentation performance, age-related associations with GM and total creatine (tCr) were examined.

**Results:** Significant differences (*p* < 0.0001) were observed in tissue fraction estimates between segmentation tools, leading to differences in metabolite concentration estimates of up to 9% under identical acquisition and modeling conditions. Although the strength of the correlation varied between segmentation methods, no statistically significant differences were found.

**Conclusion:** The choice of segmentation methodology contributed substantially to variability in MRS “absolute” metabolite concentration estimates. These results underscore the need for transparent segmentation reporting to ensure reproducibility and cross-study comparability in MRS research. Quantifying the segmentation-driven variability allows researchers to contextualize cross-study differences, helping determine whether observed effects are methodological or biologically meaningful.

## 1. Introduction

Magnetic resonance spectroscopy (MRS) is a non-invasive technique used to assess the metabolic composition of a biological sample^1^. By estimating metabolite concentrations in vivo, MRS provides insights into normal and pathological biological processes throughout the body. Its application to the brain has yielded major insights into sensory processing mechanisms^2–5^, chronic pain conditions^6,7^, neurodegenerative diseases^8,9^, psychiatric disorders^10^, and metabolic changes associated with aging^11^. For MRS to support clinical decision-making, however, it is critical to minimize methodological variability that can obscure or inflate physiological signals. Expert consensus has identified this variability, spanning data acquisition, preprocessing, and quantification, as a key barrier to the clinical translation of MRS^12^. While several factors contribute to this variability, including differences in relaxation correction^13^ and modeling approaches^14^, one critical source of variance that is understudied is tissue segmentation, which is essential for the tissue correction applied during water-referenced MRS quantification. Previous work at 1.5 T has demonstrated that variability in image segmentation approaches, particularly in partial volume estimates of cerebrospinal fluid (CSF) and gray matter (GM), can substantially impact metabolite concentration estimates^13^. For example, differences of up to 14% for *N*-acetylaspartate (NAA) have been reported depending on the segmentation method used, even when using multispectral imaging data^13^.

“Absolute” concentration estimates in biochemical units are increasingly favored due to their biological interpretability and potential for cross-study comparisons^15^. While this approach relies on several assumptions beyond the scope of the present study, such as fixed water content and relaxation properties^1,13,15,16^, these calculations depend heavily on accurate estimates of GM, white matter (WM), and CSF fractions within the MRS voxel. Accurate tissue segmentation, derived from additional anatomical MRI scans, is therefore essential for reliably estimating metabolite quantification. While GM and WM fractions inform the application of appropriate relaxation corrections, the CSF fraction has a particularly strong influence on final concentration estimates due to its assumed lack of metabolite signal^15^ and vastly different relaxation properties compared to GM and WM.

While numerous automated software packages are publicly available for brain tissue segmentation, specifically for GM, WM, and CSF, each package applies different mathematical and analytical approaches to tissue classification and, in some cases, their use of prior knowledge (e.g., from a brain template or atlas), introducing a wide range of segmentation differences^17–19^. For example, advanced normalization tools (**ANTs**) combines Markov random field (MRF) theory with template-based priors to inform tissue classification ^18^. In contrast, FMRIB software library (**FSL)** relies on hidden Markov random field (HMRF) models, where tissue class membership is determined by both voxel intensity and the contextual constraint of neighboring voxels^19^. Finally, statistical parametric mapping (**SPM)** uses a combination of a brain prior and a series of Gaussian mixture models (GMMs) to perform tissue segmentation^17^. While previous studies have reported that segmentation approaches can yield variable tissue estimates^13,20^, their direct impact on tissue- and relaxation-corrected single voxel MRS metabolite quantification has not been systematically characterized at 3 T. This study addresses this gap by evaluating how segmentation differences propagate through quantification pipelines and influence metabolite estimates.

Specifically, we assessed the impact of segmentation variability on metabolite concentration estimates using a test-retest in vivo MR dataset. By applying outputs from three commonly used segmentation tools (i.e., ANTs, FSL, and SPM) to identical MRS data, we quantified how differences in tissue fraction estimates propagate to estimated absolute metabolite concentrations. We hypothesized that these tools would yield divergent tissue classifications, which, in turn, would introduce significant variability in MRS-derived absolute metabolite levels. Additionally, age-related associations in both normalized GM fraction and metabolite concentrations (tCr) were used as biological benchmarks to evaluate which segmentation approach most accurately reflects known neuroanatomical and neurochemical relationships with age^21–25^. This strategy aligns with prior work that validated MR-based measures by correlating them with well-characterized biological variables^26,27^.

## 2. Methods

### 2.1 Experimental details

Ethical approval for this study was obtained from the Weill Cornell Medicine Institutional Review Board (protocol #0807009883), and all participants provided written informed consent before their participation. Sixteen healthy adults (6 males, 10 females; mean age ± SD: 38.4 ± 18.2 range: 19–66 years) were enrolled in a separate study focused on test-retest reliability^28^. Each participant completed two MR scan sessions, with a median interval of 0 days between sessions (range: 0–29 days), as most were scanned on the same day or within days, with a single participant contributing the longest interval. The exclusion criteria were for individuals with contraindications to MRI or a history of neurological or psychiatric disorders. The study design, including MRI/MRS data acquisition and segmentation workflow, is illustrated in **Figure 1**.

**Figure 1.**
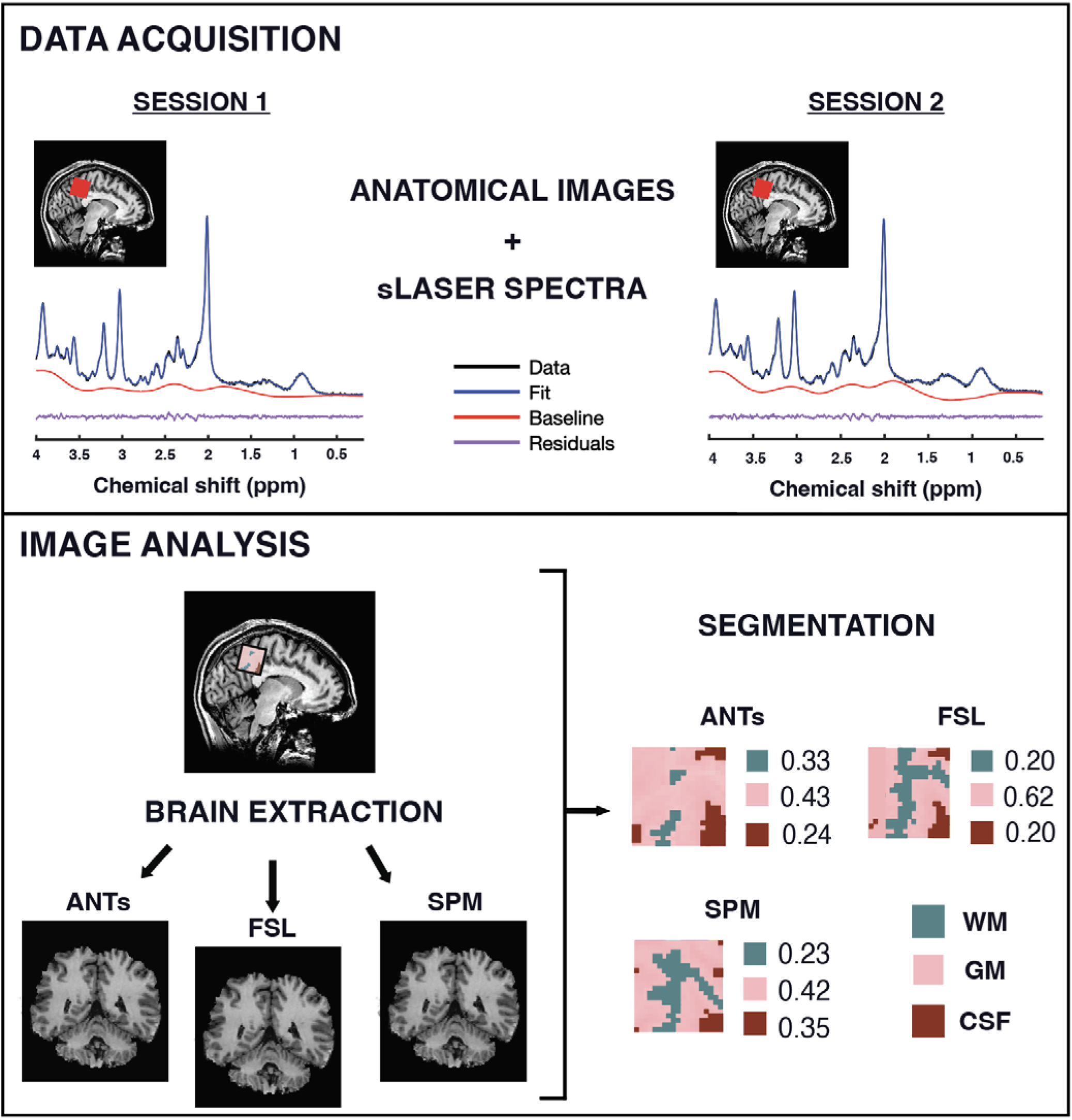
Study design of MR data acquisition and analysis. A 27-mL voxel was positioned in the medial parietal lobe to localize MRS data. A representative MRS spectrum from one participant across two scan sessions, including the raw spectral data (black), LCModel fit (blue), baseline estimate (red), and residuals (purple). Brain extraction and segmentation were performed using three software tools: ANTs, FSL, and SPM.

**Figure 2.**
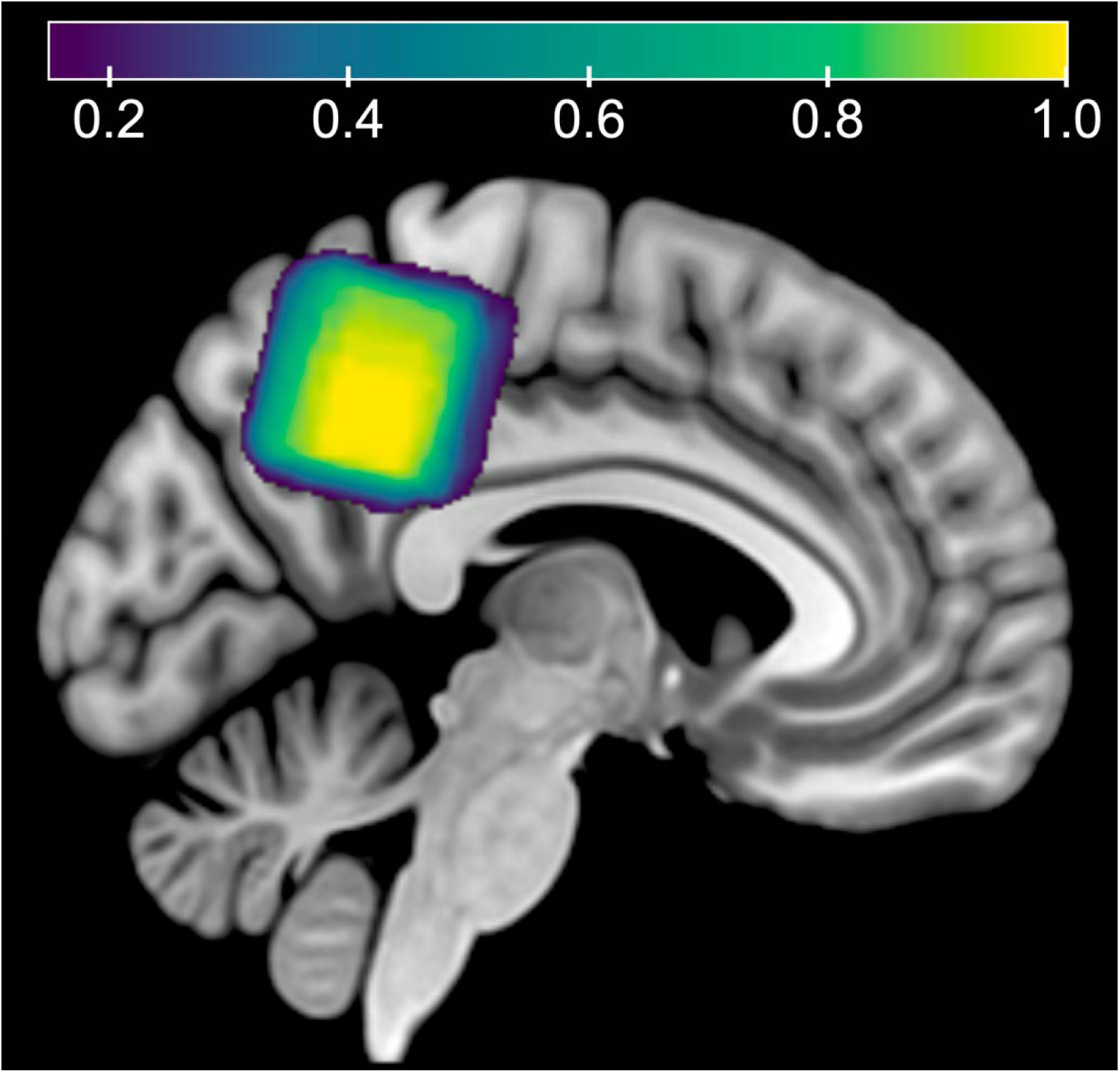
Voxel placement map. Showing the average overlap between scan sessions 1 and 2 in the medial parietal lobe for all participants in MNI152 template space. The color bar denotes the estimated overlap, where 1.0 equates to 100% overlap.

### 2.2 MR Scanning Protocol

Data were collected on a General Electric (GE) Discovery 3T MR750 scanner using a ^1^H 32-channel phased-array RF head coil for receive and a body coil for transmit. High-resolution 3D T_1-_weighted Brain Volume Imaging (BRAVO) structural scans (fast spoiled gradient echo (FSPGR); TR/TE/TI = 12.2/5.2/725 ms; flip angle = 7°; voxel resolution = 0.9 × 0.9 × 1.5 mm^3^; matrix size = 256 × 256; slices = 124; parallel acceleration factor = 2) were first acquired for accurate voxel placement in each scan session. Single-voxel MRS data were acquired using a semi-localization adiabatic selective refocusing (sLASER)^29^ sequence with a TR/TE of 2000/35 ms, a spectral width of 5000 Hz, 4096 data points, and 64 transients (**Figure 1)**. The voxel resolution was 3 × 3 × 3 cm³, and the MRS voxel was placed in the medial parietal lobe (**Figure 3**). Water suppression was performed using variable power RF pulses with optimized relaxation delays (VAPOR)^30^. All sequence details for the MRS acquisition and analysis can also be found in the MRSinMRS table in the supplemental materials^12^.

**Figure 3.**
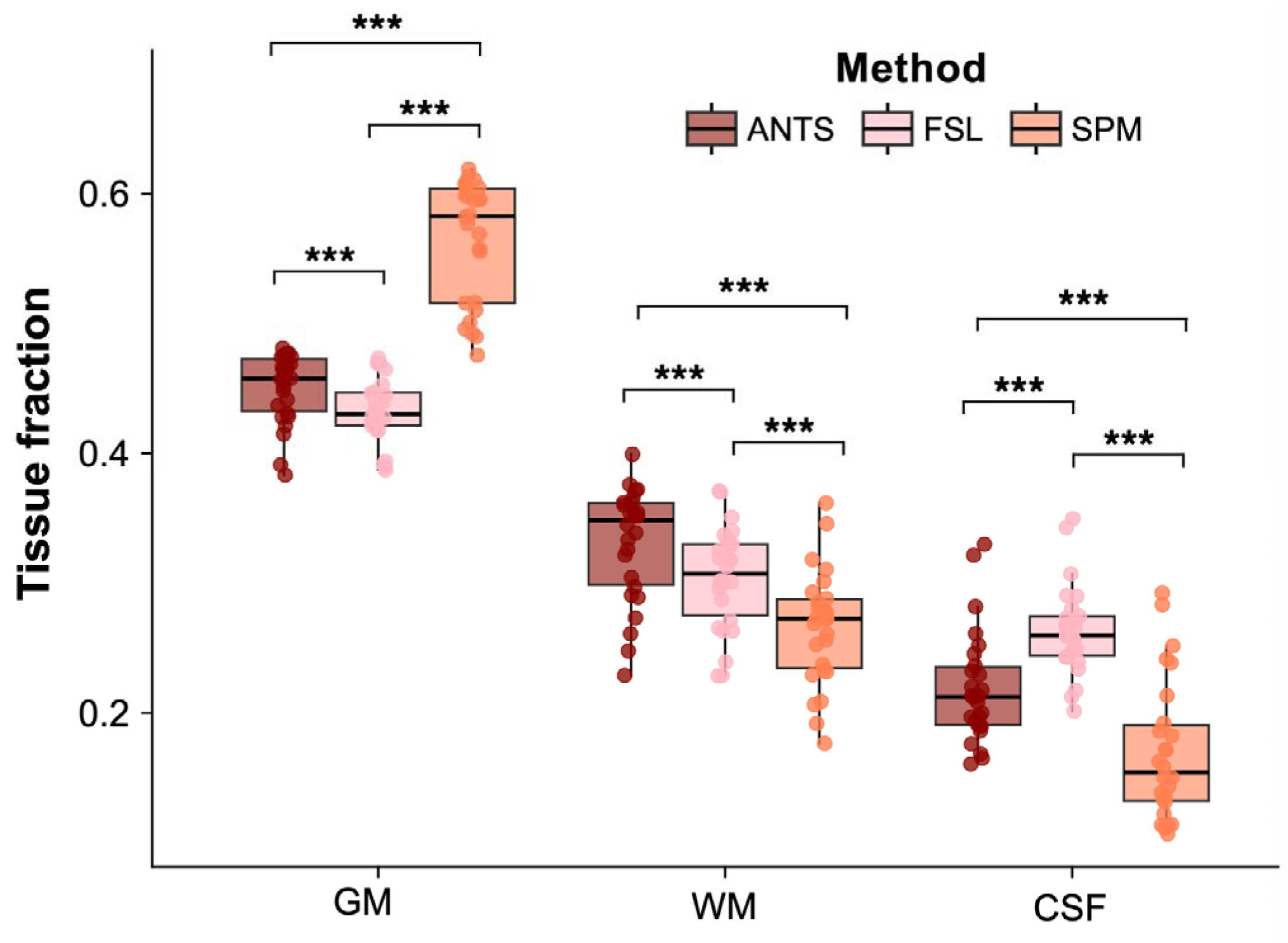
Comparison of tissue segmentation output across methods. Gray matter (GM), white matter (WM), and cerebrospinal fluid (CSF) fractions are shown on the *x*-axis. Data were pooled over both sessions (*n* = 26). *p* < 0.05 = *, *p* < 0.01 = **, *p* < 0.001 = ***

### 2.3 MR Image Segmentation

The 3D T_1_-weighted structural data were brain-extracted and segmented into GM, WM, and CSF using each of the following software packages: ANTs, FSL, and SPM. Specifically, brain extraction was performed separately within each segmentation software. This approach ensured that the brain extraction and segmentation steps were consistent with the design of each software. To ensure comparability across methods, tissue fraction maps were verified to sum to 1 within each MRS voxel and were normalized if this condition was not already met.

Further detail on each segmentation approach is described below.

#### 2.3.1 ANTs (Advanced Normalization Tools) Atropos

ANTs Atropos (v2.5.1) employs a Bayesian framework coupled with a non-parametric finite mixture model (FMM), which can switch between a Gaussian mixture model (GMM) or an FMM depending on the assumed distributions of each tissue class, to optimize the voxel classification into specific tissue classes. The FMM assumes voxel-wise independence when estimating the likelihood of a voxel belonging to a specific class, based on observed intensity across the image. To incorporate spatial coherence, Atropos integrates prior probabilities, modeled using either markov random fields (MRFs) or template-based labeled priors, selected by the user. The soft expectation-maximization (sEM) algorithm is then used to iteratively find the optimal voxel classifications by maximizing the posterior probability and redefining the mixing parameter, gamma-k, at each iteration. This balances the contribution of the likelihood (from the FMM or GMM) with the prior probabilities and allows voxels to be assigned to multiple classes. For consistency in comparison, we used the same priors in ANTs as those applied in SPM12.

#### 2.3.2 FSL (BET & FAST) FMRIB’s Automated Segmentation Tool (FAST)

FAST (v6.0.5)^19^ employs a stochastic approach by using the hidden Markov random field with expectation maximization (HMRF-EM) and the iterative conditional modes (ICM) to perform three tissue segmentations. This algorithm incorporates spatial priors, as a voxel’s membership in a specific tissue class depends on the influence of the surrounding voxels. This HMRF-EM framework employs a maximum a posteriori estimate to calculate estimates of the bias field and class labels, while maximum likelihood is used to estimate the model parameters.

#### 2.3.3 Statistical Parametric Mapping (SPM)

The SPM12 (V7.2.1.9) unified segmentation algorithm uses a generative model that jointly performs tissue classification, intensity bias correction, and image registration within a probabilistic Bayesian framework. The probability of a voxel belonging to a specific class is modeled by a GMM, where the likelihood function^17^ assumes independence. Similar to Atropos, voxel spatial coherence is incorporated via spatial priors. The algorithm applies intensity non-uniformity bias correction to log-transformed data, also within a GMM framework. Finally, to capture voxel spatial dependence a modified version of the International Consortium for Brain Mapping (ICBM) Tissue Probabilistic Atlas as the spatial prior is used, where a GMM approach is still used; however, each tissue is represented by its own set of Gaussian distributions: GM = 3, WM = 2, CSF = 1, parameterizing tissue classifications through a combination of Gaussians that represent a specific tissue type. A regularization term is added to the model to penalize nonuniformity or the presence of an intensity bias field. Traditional EM is employed to iteratively estimate the parameters of the probabilistic model, which characterizes the underlying tissue classifications and biases present in the image data. Optimizing the objective function and incorporating the regularization term, making this a parametric approach.

### 2.4 MRS Data Processing

MRS data were processed in Osprey (v 2.9.5)^31^ and fitted using the embedded LCModel wrapper (v6.3-1N). A custom sLASER basis set was generated using FID-A^32^ with the following simulation parameters: twenty metabolites (alanine (Ala), ascorbate (Asc), aspartate (Asp), creatine (Cr), gamma-Aminobutyric Acid (GABA), glucose (Glc), glutamine (Gln), glutamate (Glu), glycine (Gly), glycerophosphocholine (GPC), glutathione (GSH), myo-Inositol (mI), lactate (Lac), *N*-acetylaspartate (NAA), *N*-acetylaspartylglutamate (NAAG), phosphocholine (PCh), phosphocreatine (PCr), phosphoethanolamine (PE), scyllo-inositol (scyllo), and taurine (tau)), a line width (LW) of 2 Hz, a spectral width of 5000 Hz, and 64 × 64 spatial grid points^33^ (*see Code Availability*). The metabolite and water signals were then used to calculate the molar concentration of tNAA, tCr, tCho, mI, Glu, and the combination of Glu and Gln (Glx)^15^(*see Code Availability*). We report results for tCr in the main text; results for the remaining metabolites are presented in the Supplementary Material. This choice was made because segmentation-related variability propagates similarly across all metabolites.

### 2.5 MRS Quantification

Anatomical segmentation variability was evaluated across three approaches, ANTs, FSL, and SPM, by extracting fractional volumes of GM, WM, and CSF from the MRS voxel. The varying tissue fractions were used to compute [M]_molar_ according to Gasparovic et al.^15,34^

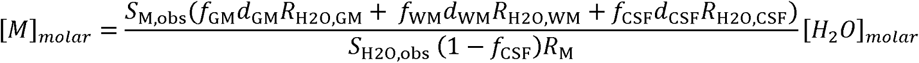

Where:

- *S*_M,obs_ and *S*_H2O,obs_ are the observed metabolite and water signals (as modeled by Osprey), *f_x_* is the fractional volume of GM, WM, and CSF within the voxel;
- *d_x_* is the density of water in GM, WM, and CSF (*d*_GM_ = 0.78, *d*_WM_ = 0.65, and *d*_CSF_ = 0.97) ^15,16^;
- *R*_H2O,*x*_ is the relaxation attenuation factor for water in GM, WM, and CSF;
- *R*_M,_ is a scaling factor that accounts for relaxation times of metabolite protons averaged over GM and WM^34,35^.
- [H_2_O] is the molar concentration of pure water (55.51 mol/L^15^).

Relaxation attenuation terms were calculated using a standard exponential model^15,34^, where water-specific *T*_1_ and *T*_2_ values were taken from literature for GM, WM, and CSF^36^. For metabolites, we used mean T_1_ and T_2_ values from prior studies across GM and WM^35,37^ (**see Supplementary Material**). Note that the number of protons for each metabolite is encoded in the basis set (see Code Availability).

### 2.6 Statistical Analysis

The statistical analysis was performed using R (4.4.0). A significance level of p < 0.05 was used for all inference tests. To test if metabolite concentration values were normally distributed, Shapiro–Wilk tests were performed for each metabolite (tNAA, tCr, mI, Glu and Glx) within each segmentation dataset for each session with Bonferroni correction for multiple comparisons.

#### 2.6.1 Between-Session Differences

A paired t-test was used to assess the differences between the two scan sessions, after determining the normality of the data. Quality metrics, including the Cr signal-to-noise ratio (Cr SNR), water peak full width at half maximum (H_₂_O FWHM), and the fit quality index (defined as the ratio of the residual sum of squares to the squared standard deviation of the noise), were evaluated for session-related differences. Bonferroni corrections were applied to account for testing these outcome measures.

#### 2.6.2 Repeated-Measures Analysis Using Linear Mixed-Effects Models

a. To evaluate the effects of segmentation method on tissue composition estimates, linear mixed-effects models were fitted separately for each tissue type (GM, WM, CSF) using the lme function in R^38^ as follows: (Fraction ∼ Method * Session, random = ∼1|Participant,data). Each model included segmentation method (ANTs, FSL, SPM), session, and their interaction as fixed effects, with participant modeled as a random intercept to account for repeated measures. The significance of fixed effects was assessed using ANOVA-style tables derived from the models.
b. To assess whether segmentation method and session influence MRS metabolite quantification, linear mixed-effects models were similarly fitted separately for each metabolite (e.g., tNAA, tCr, Glu, Glx, mI), with segmentation method, session, and their interaction as fixed effects, and participant as a random intercept. The model was specified as: lme (Metabolite ∼ Method * Session, random = ∼1|Participant,data). ANOVA-style tables summarizing the fixed effects were generated for each model. Post hoc pairwise comparisons were performed using Tukey’s HSD.

#### 2.6.3 Correlational Analysis

Pearson’s correlation coefficients were computed to examine the relationship between participant age and normalized GM tissue fraction (fGM / [fGM + fWM]) derived from each segmentation method (e.g., ANTs, FSL, SPM). Correlation strength (*r*), coefficients of determination (*r*^2^) (i.e., effect sizes), and associated *p*-values are reported. To statistically compare dependent correlations sharing a common variable (e.g., age) across segmentation methods, Steiger’s *Z*-test for comparing correlated correlation coefficients was performed^39^. This test accounts for the dependency between correlations derived from the same sample.

#### 2.6.4 Percent Differences

Percent differences were calculated to summarize the relative differences in metabolite concentration between scan sessions (within-method comparisons) and between segmentation methods (between-method comparisons). First, within-method variability was evaluated by calculating the percent change in metabolite concentration between the two scan sessions for each participant, separately for each segmentation method. The mean percent change across participants was then computed for each metabolite and segmentation method. Second, to assess between-method variability, the average concentration across sessions was calculated for each metabolite, participant, and segmentation method. Percent differences between methods were then determined by comparing the mean concentrations from each pair of segmentation methods (SPM vs. FSL, SPM vs. ANTs, and FSL vs. ANTs).

## 3. Results

Data from two participants were excluded due to a lack of relevant consent for data sharing, and data from one participant were excluded due to the participant wishing to end the scan. For the remaining cohort (*n* = 13), between-session differences were evaluated. MRS quality metrics for both sessions are summarized in **Table 1**. Shapiro-Wilk tests indicated no significant departures from normality in either session (all p > 0.05). Full results are provided in the supplementary material, with density plots included.

**Table 1.** MRS Spectral Quality Outcome Measures.

|  | CR SNR | H<sub>2</sub>O FWHM | FIT QUALITY |
| --- | --- | --- | --- |
| SESSION 1 | 152.43 ± 14.14 | 7.27 ± 0.66 | 2.36 ± 0.29 |
| SESSION 2 | 156.03 ±13.49 | 7.23 ± 0.63 | 2.67 ± 0.59 |

### 3.1 Metabolite Quality Metrics Between Sessions

No significant session-to-session differences in spectral quality metrics (Cr SNR: t = -0.98, p= 0.99; H_2_O FWHM: t = 0.26, p < 0.99; fit quality index: t= - 2.07, p = 0.18) were found.

### 3.2 Effect of Segmentation Method on Tissue Fraction Estimates

For GM tissue fraction estimates, there was a significant main effect of segmentation method (F(2,60) = 434.23, p < 0.0001), with no significant effect of session (F(1,60) = 0.54, p = 0.47) and no significant interaction (F(2,60) = 0.05, p = 0.95). Estimated marginal means averaged over sessions showed that SPM produced the highest GM fraction (mean = 0.56, 95% CI [0.54, 0.58]), followed by ANTs (mean = 0.45, 95% CI [0.43, 0.47]) and FSL (mean = 0.43, 95% CI [0.41, 0.45]). Post-hoc Tukey-adjusted pairwise comparisons for segmentation method revealed that ANTs estimates were significantly higher than FSL (mean difference = 0.02, SE = 0.01, p = 0.0018), and SPM was significantly higher than both ANTs (mean difference = 0.11, SE = 0.01, p < 0.0001) and FSL (mean difference = 0.13, SE = 0.01, p < 0.0001). Session means were similar (session 1: mean = 0.48, 95% CI [0.46, 0.50]; session 2: mean = 0.48, 95% CI [0.46, 0.50]), with no significant difference between sessions (mean difference = 0.00, SE = 0.01, p = 0.47) (**Figure 3**).

For WM tissue fraction estimates, there was a significant main effect of segmentation method (F(2,60) = 120.54, p < 0.0001), with no significant effect of session (F(1,60) = 0.98, p = 0.33) and no significant interaction (F(2,60) = 0.24, p = 0.79). Estimated marginal means averaged over sessions showed that ANTs produced the highest WM fraction (mean = 0.33, 95% CI [0.31, 0.36]), followed by FSL (mean = 0.30, 95% CI [0.28, 0.33]) and SPM (mean = 0.27, 95% CI [0.24, 0.29]). Post-hoc Tukey-adjusted pairwise comparisons for segmentation method revealed that ANTs estimates were significantly higher than both FSL (mean difference = 0.03, SE = 0.00, p < 0.0001) and SPM (mean difference = 0.06, SE = 0.00, p < 0.0001), and FSL estimates were significantly higher than SPM (mean difference = 0.04, SE = 0.00, p < 0.0001). Session means were similar (session 1: mean = 0.30, 95% CI [0.28, 0.32]; session 2: mean = 0.30, 95% CI [0.28, 0.33]), with no significant difference between sessions (mean difference = -0.00, SE = 0.00, p = 0.33) (**Figure 3**).

For CSF tissue fraction estimates, there was a significant main effect of segmentation method (F(2,60) = 185.62, p < 0.0001), with no significant effect of session (F(1,60) = 0.01, p = 0.90) and no significant interaction (F(2,60) = 0.08, p = 0.93). Estimated marginal means averaged over sessions showed that FSL produced the highest CSF fraction (mean = 0.26, 95% CI [0.24, 0.29]), followed by ANTs (mean = 0.22, 95% CI [0.19, 0.25]) and SPM (mean = 0.17, 95% CI [0.14, 0.20]). Post-hoc Tukey-adjusted pairwise comparisons for segmentation method revealed that ANTs estimates were significantly lower than FSL (mean difference = - 0.04, SE = 0.00, p < 0.0001), while ANTs estimates were significantly higher than SPM (mean difference = 0.05, SE = 0.00, p < 0.0001), and FSL estimates were significantly higher than SPM (mean difference = 0.09, SE = 0.00, p < 0.0001). Session means were nearly identical (session 1: mean = 0.22, 95% CI [0.19, 0.24]; session 2: mean = 0.22, 95% CI [0.19, 0.25]), with no significant difference between sessions (mean difference = -0.00, SE = 0.01, p = 0.90) (**Figure 3**).

### 3.3 Effect of Segmentation Method on Quantified Metabolites

**For tCr**, the analysis showed a significant main effect of segmentation method, F(2, 60) = 64.63, p < 0.0001, with no significant session effect, F(1, 60) = 0.59, p = 0.44, and no interaction (of session and segmentation method), F(2, 60) = 0.06, p = 0.94 (ANTS: mean = 7.19, 95% CI [7.01, 7.37]; FSL: mean = 7.59, 95% CI [7.40, 7.77]; SPM: mean = 6.89, 95% CI [6.71, 7.07]).All post-hoc pairwise comparisons between segmentation methods are provided in **Table 2 and Figure 4**. Results for additional metabolites are summarized in **Table S4** (Supplementary Material).

**Figure 4.**
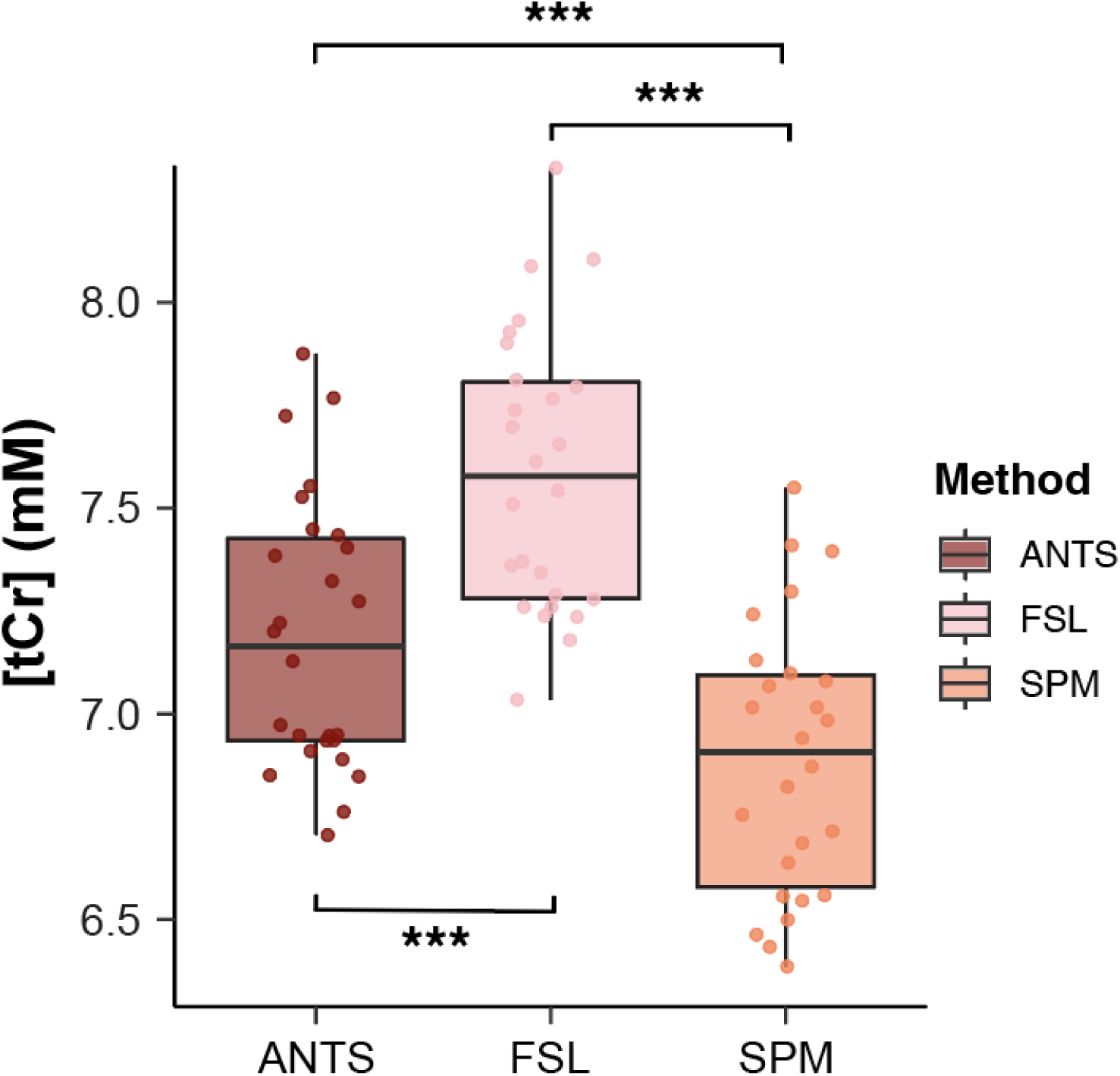
Estimated concentrations (mM) of tCr across the three segmentation datasets (ANTs, FSL, and SPM). Results are shown from a repeated-measures ANOVA with Tukey post hoc-corrected pairwise comparisons. Significant differences in metabolite concentration between datasets are indicated by asterisks (*p* < 0.05); tCr, total creatine. *p* < 0.05 = *, *p* < 0.01 = **, *p* < 0.001 = ***

**Table 2.** Tukey-Adjusted Pairwise Comparisons of tCr levels Across Segmentation Methods.

| METHOD | ESTIMATE | SE | DF | T-RATIO | P-VALUE |
| --- | --- | --- | --- | --- | --- |
| ANTS – FSL | -0.40 | 0.06 | 60 | -6.48 | < 0.0001 |
| ANTS – SPM | 0.30 | 0.06 | 60 | 4.85 | < 0.0001 |
| FSL – SPM | 0.70 | 0.06 | 60 | 11.33 | < 0.0001 |

Mean percentage differences in tCr levels across segmentation methods and scan sessions are presented in **Figure 5**. Within-software differences between sessions were substantially smaller than those observed between segmentation programs. The largest difference was observed between FSL and SPM (∼9%). This pattern was similarly observed across other metabolites; full results for all metabolite comparisons are provided in **Figure S2**.

**Figure 5.**
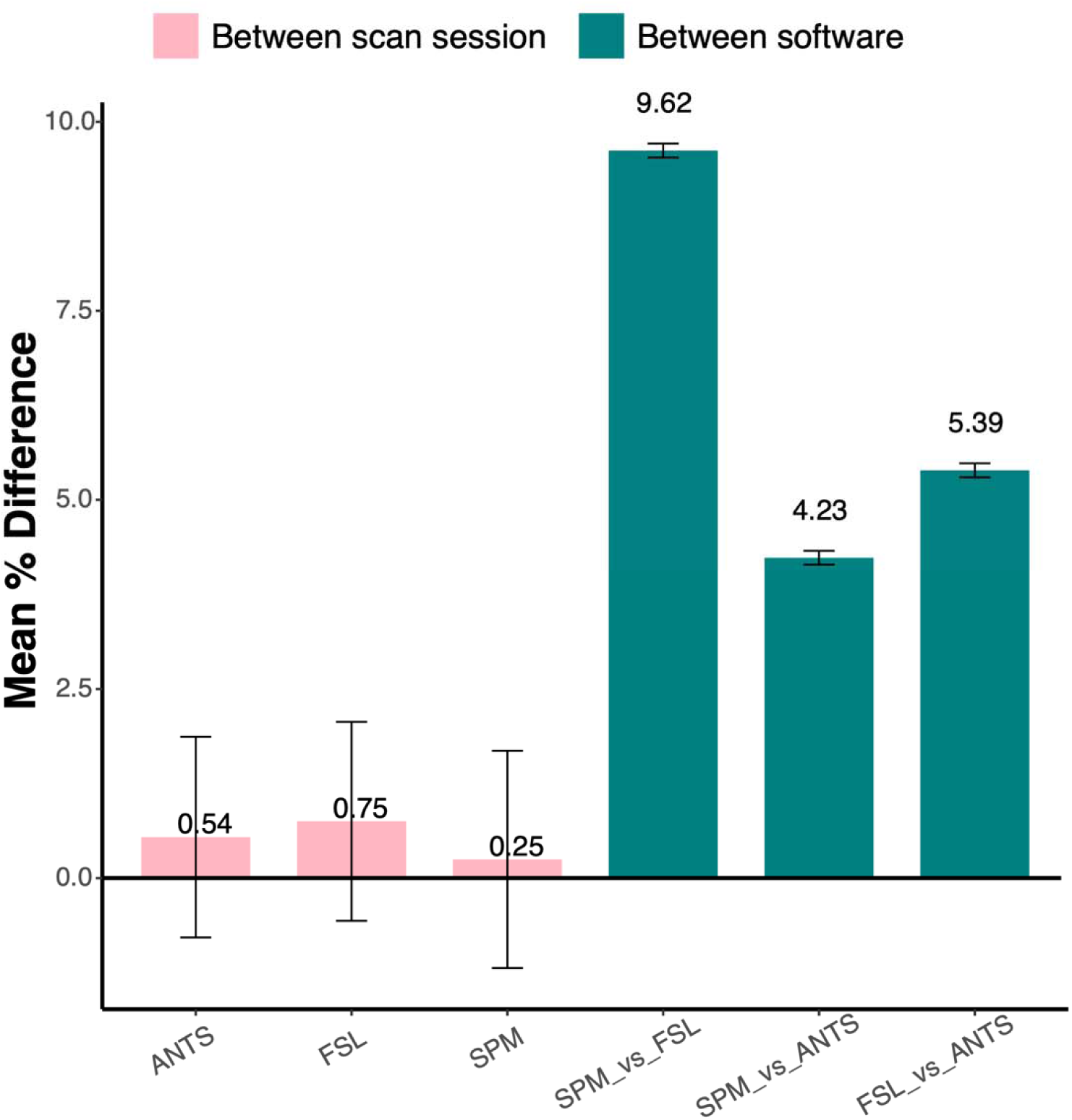
Mean percentage differences for tCr levels across methods. Bars represent the mean percentage difference between scan sessions (light pink) and between software packages (teal). Error bars denote the standard deviation. tCr, total creatine.

### 3.4 Assessment of Segmentation Methods Using Age-Related Biological Trends

Age was negatively associated with normalized GM fraction for all segmentation tools: ANTs [r = -0.72, r² = 0.51, p = 0.01], FSL [r = -0.66, r² = 0.44, p = 0.03], and SPM [r = -0.69, r² = 0.48, p = 0.01]. Age was positively associated with tCr, with the strongest association observed for SPM [r = 0.62, r² = 0.38, p = 0.02], followed by ANTs [r = 0.32, r² = 0.10, p = 0.29] and FSL [r = 0.22, r² = 0.05, p = 0.47] (**Figure 6)**. Steiger’s Z-tests revealed no significant differences in the strength of correlations across segmentation methods. For normalized GM ratio and age, comparisons yielded: SPM vs FSL (z = –0.39, p = 0.70), SPM vs ANTs (z = –0.39, p = 0.70), and FSL vs ANTs (z = –0.39, p = 0.70). Similarly, for age–tCr associations: SPM vs FSL (z = – 0.83, p = 0.40), SPM vs ANTs (z = –0.83, p = 0.40), and FSL vs ANTs (z = –0.83, p = 0.40).

**Figure 6.**
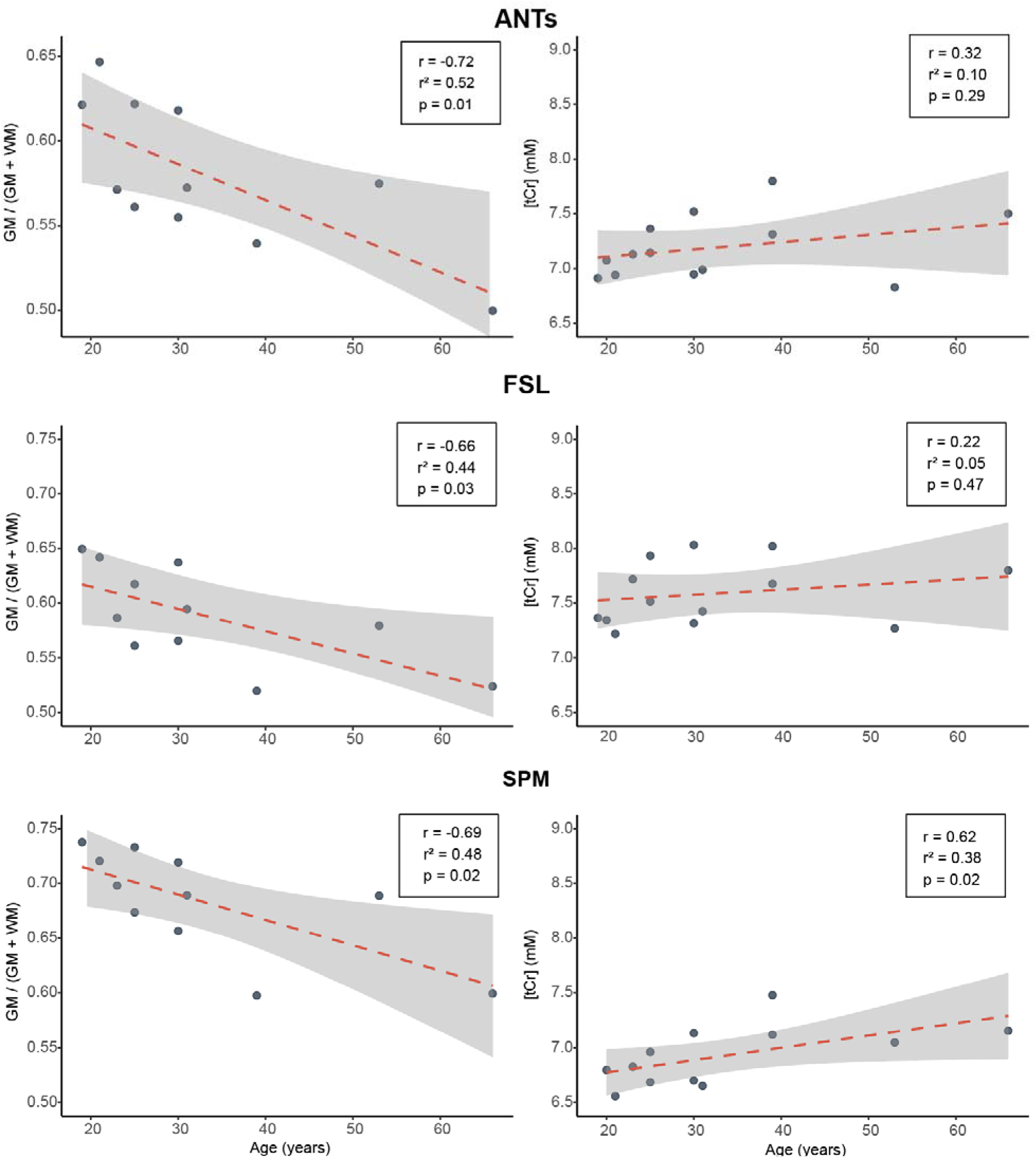
Age-related changes in GM and tCr. Plots on the right column show the association between mean normalized gray matter fraction across sessions and participant age, as calculated by three segmentation programs: ANTs, FSL, and SPM. On the left column, the mean tCr across sessions is plotted against participant age, as calculated by the three segmentation programs. Each plot displays the Pearson correlation coefficient (*r*), coefficient of determination (*r*²), and corresponding p-value, illustrating the strength and significance of the relationship (*n* = 26).

## 4. Discussion

This study demonstrates that metabolite concentration estimates derived from single-voxel MRS are substantially influenced by the choice of tissue segmentation method, even under identical acquisition and modelling conditions. Although metabolite levels remained stable across scan sessions, supporting consistent data acquisition, considerable differences emerged in absolute metabolite estimates when different tissue segmentation methods were used to calculate tissue fractions. tCr, like other commonly measured MRS metabolites presented in the supplementary material, plays a central role as a clinical and research marker, particularly in studies of schizophrenia^40^, neurodegeneration^41–44^, and cancer^45^. The greatest difference was observed between SPM and FSL, with estimates varying by as much as ∼ 9%, a considerable discrepancy, especially considering that many clinical and sensory MRS studies aim to detect subtle changes or similar or smaller magnitude^46–48^. Moreover, variability in segmentation pipelines across studies may limit the comparability of published normative ranges and thresholds for pathology, thereby complicating efforts in meta-analyses and hindering translational application.

This study highlights the influence of segmentation methodology on estimated absolute concentrations, given that many contemporary MRS processing pipelines leverage established neuroimaging processing tools like SPM or FSL^31,49^. These findings have important implications for different study designs in MRS research. Within-participant comparisons (e.g., task versus rest, pre-versus post-intervention) are likely to be the least affected by segmentation variability, as tissue misclassification errors may cancel out when anatomical structures and voxel placement are consistent across conditions. In contrast, within-study, between-participant comparisons, such as group differences, may be moderately impacted, especially in cohorts with greater anatomical variability due to aging or pathology. In these cases, systematic biases in tissue proportion estimates as a function of anatomy could distort group-level inferences. Longitudinal studies, particularly those spanning developmental or degenerative changes, may also be susceptible, as shifts in anatomy over time could interact with segmentation biases in complex ways. The greatest vulnerability arises in across-study comparisons attempting to establish estimated absolute concentrations as the primary outcome, for example, quantitative biomarker thresholds. Here, the choice of segmentation tool can produce up to several millimolar differences in metabolite estimates even when acquisition, processing, modeling and relaxation correction parameters are otherwise identical, posing significant challenges for data harmonization, meta-analyses, and clinical translation.

Metabolite concentration estimates rely on assumed values for tissue-specific relaxation times and water content, often derived from literature averages that may not reflect individual physiology. Prior work has shown that metabolite relaxation times differ by tissue type and vary with acquisition parameters, yet these parameters are seldom measured directly in vivo due to time constraints. While this limitation is not the focus of the present study, it represents a fundamental constraint on the precision of estimated absolute quantification, further compounding the variability introduced by segmentation. Tissue segmentation plays a particularly important role in estimating absolute concentrations, as tissue fractions directly inform relaxation corrections. Each tissue compartment has distinct water content and relaxation properties. Among these, CSF is particularly challenging: not only are its relaxation parameters less consistently reported in the literature^50^, but its volume increases with age-related atrophy^51^. CSF relaxation times differ more strongly from GM and WM relaxation times than GM and WM relaxation times differ from each other. Inaccurate estimation of CSF fractions can thus introduce significant bias in the correction of the water signal and thus have a major influence on metabolite estimates. Beyond these considerations, Maddock emphasizes that even when using water-referenced metabolite concentration estimates, they will still be confounded by shared sources of nuisance, a concern later confirmed by Prisciandaro et al.^52,53^ In particular, Maddock highlights segmentation-related noise as a key contributor, noting that variability in voxel-wise tissue fraction estimates, used for relaxation and water-content correction, can introduce shared error across metabolites within the same voxel, leading to spurious correlations even in the absence of true biological associations^52^. Furthermore, estimated absolute quantification into molar concentrations is not the only instance in which accurate tissue segmentation is critical. Accurate segmentation is also necessary for reliable partial volume correction, as well as for other quantification approaches, such as metabolite ratios. Finally, we note that there are also tissue-dependent differences in metabolite relaxation times and in intrinsic metabolite concentrations (i.e., metabolites are inherently concentrated differently across tissue types). Such corrections have been considered before^54–56^, but were not applied in this study to avoid over-complicating the scope of our investigation.

Beyond statistical agreement, segmentation methods can also be evaluated by how well they align with known biological patterns^21,23,24^. In the absence of a ground truth, such biologically plausible trends offer a complementary form of validation. All segmentation tools showed significant associations between age and GM fraction, consistent with well-established age-related tissue changes^57^, although these differences were not statistically significant from each other; this may reflect method-specific sensitivity to age-related morphological changes. For tCr, SPM was the only method to show an association with age exceeding a Pearson correlation coefficient of 0.4. While our goal was not to establish definitive age–metabolite relationships, this correlation provides a useful validity check: no method produced implausible or reversed patterns, and the observed associations were in line with prior studies reporting a positive relationship between Cr levels and age using single-voxel MRS ^21^, MRSI ^24^, and EPSI^25^. Notably, the single-voxel study also employed SPM for tissue segmentation, raising the possibility that the choice of segmentation may partly explain this alignment. However, replication with other acquisition and segmentation strategies suggests a potentially robust biological signal rather than a tool-specific artifact. Importantly, the age–tCr relationship is less well characterized than the well-documented association between age and GM atrophy^57,58^, warranting cautious interpretation and further study. Our sample size limits the ability to detect small differences, but these correlations serve to demonstrate due diligence and provide context for the relative plausibility of segmentation outputs. While manual segmentation is often considered the gold standard, it too has limitations, such as rater bias, making biologically anchored metrics essential for benchmarking. To complement these findings, future work should develop analytical frameworks for evaluating segmentation methods. Mathematical modeling approaches, such as uncertainty and sensitivity analyses, could help quantify how segmentation variability propagates through to MRS quantification outcomes under controlled conditions^59^.

A key limitation of this study is the relatively small sample size, which may limit the generalizability of the findings and reduce the statistical power to detect subtle differences between segmentation methods. Additionally, the absence of a definitive ground truth tissue segmentation makes it challenging to determine absolute accuracy across tools, even as manual segmentations may be subjective in nature which introduces its own set of biases. Notably, all analyses were conducted in a healthy population; thus, it remains unclear how well these segmentation algorithms perform in the context of neurological or structural abnormalities, which may further constrain the broader applicability of the results. Although we matched prior probability maps across methods to ensure comparability, differences in algorithmic implementation may still have influenced the results. Future work will investigate how the selection and tuning of prior probabilities impact segmentation outcomes and aim to establish broader biological and methodological benchmarks.

## 5. Conclusions

The impact of segmentation methods from three widely used software tools was assessed to determine their effect on metabolite estimates using a test-retest MR dataset. Significant differences were observed in the absolute quantification estimates of tCr, independent of session effects. These findings highlight the importance of segmentation software choice in MRS quantification, especially when comparing data across studies that have used different segmentation methods. Understanding the extent of this variability is essential not only for improving the reliability and reproducibility of MRS studies but also for advancing our knowledge of brain metabolism in health and disease.

## Code Availability

All MRS data and code used in this study are publicly available on OpenNeuro. [doi:10.18112/openneuro.ds006444.v1.0.0].The basis set generation code is available on GitHub at: https://github.com/arcj-hub/BasisSetSimulation. All anatomical segmentation scripts are openly available on GitHub at: https://github.com/arcj-hub/BIASS-INVIVO. Statistical analysis code and the molar quantification are included in the OpenNeuro repository.

## Funding

MM is supported by National Institutes of Health grant K99EB028828. CCF is supported by NIH DP2NS127704. GO and the Osprey software development are supported by NIH R01EB035529 and R21EB033516. GO is a paid consultant for Neurona Therapeutics Inc (unrelated to this work). HJZ is supported by NIH K99AG080084.

## Notes

https://openneuro.org/datasets/ds006444/versions/1.0.0

### Competing Interest Statement

The authors have declared no competing interest.

## REFERENCES

1. De Graaf, R. A. . *In Vivo NMR Spectroscopy : Principles and Techniques*. (John Wiley & Sons, Inc., 2019).

2. Betina Ip, I., et al. Combined fMRI-MRS acquires simultaneous glutamate and BOLD-fMRI signals in the human brain. Neuroimage 155, 113–119 (2017).

3. Mullins, P. G., et al. Current practice in the use of MEGA-PRESS spectroscopy for the detection of GABA. NeuroImage vol. 86 43–52 Preprint at 10.1016/j.neuroimage.2012.12.004 (2014).

4. Archibald, J., Warner, F. M., Ortiz, O., Todd, M. & Jutzeler, C. R. NEURO FORUM Sensory Processing Recent advances in objectifying pain using neuroimaging techniques. J Neurophysiol 120, 387–390 (2018).

5. Archibald, J. et al. Excitatory and inhibitory responses in the brain to experimental pain: A systematic review of MR spectroscopy studies. Neuroimage 215, (2020).

6. Foerster, B. R. et al. Reduced insular γ-aminobutyric acid in fibromyalgia. Arthritis Rheum 64, 579–583 (2012).

7. Zhao, X., Xu, M., Jorgenson, K. & Kong, J. Neurochemical changes in patients with chronic low back pain detected by proton magnetic resonance spectroscopy: A systematic review. NeuroImage: Clinical vol. 13 33–38 Preprint at 10.1016/j.nicl.2016.11.006 (2017).

8. O□z, G. Magnetic Resonance Spectroscopy of Degenerative Brain Diseases. . (: Springer., Switzerland, 2016).

9. Öz, G., et al. Clinical proton MR spectroscopy in central nervous system disorders. Radiology vol. 270 658–679 Preprint at 10.1148/radiol.13130531 (2014).

10. Richard J Maddock, M. H. B. MR spectroscopic studies of the brain in psychiatric disorders. Curr Top Behav Neurosci . (2012).

11. Haga, K. K., Khor, Y. P., Farrall, A. & Wardlaw, J. M. A systematic review of brain metabolite changes, measured with 1H magnetic resonance spectroscopy, in healthy aging. Neurobiology of Aging vol. 30 353–363 Preprint at 10.1016/j.neurobiolaging.2007.07.005 (2009).

12. Lin, A., et al. Minimum Reporting Standards for in vivo Magnetic Resonance Spectroscopy (MRSinMRS): Experts’ consensus recommendations. NMR Biomed 34, (2021).

13. Gasparovic, C. et al. Use of tissue water as a concentration reference for proton spectroscopic imaging. Magn Reson Med 55, 1219–1226 (2006).

14. Zöllner, H. J. et al. Comparison of linear combination modeling strategies for edited magnetic resonance spectroscopy at 3 T. NMR Biomed 35, (2022).

15. Near, J., et al. Preprocessing, analysis and quantification in single-voxel magnetic resonance spectroscopy: experts’ consensus recommendations. NMR in Biomedicine vol. 34 Preprint at 10.1002/nbm.4257 (2021).

16. T. Ernst, R. K. B. D. R. A. Q. of W. and M. in the H. Brain. I. C. and W. Absolute Quantitation of Water and Metabolites in the Human Brain. I. Compartments and Water. Journal of Magnetic Resonance (1993).

17. Ashburner, J. & Friston, K. J. Unified segmentation. Neuroimage 26, 839–851 (2005).

18. Avants, B. B., Tustison, N. J., Wu, J., Cook, P. A. & Gee, J. C. An open source multivariate framework for N-tissue segmentation with evaluation on public data. Neuroinformatics 9, 381–400 (2011).

19. Zhang, Y., Brady, M. & Smith, S. Segmentation of Brain MR Images Through a Hidden Markov Random Field Model and the Expectation-Maximization Algorithm. IEEE TRANSACTIONS ON MEDICAL IMAGING vol. 20 (2001).

20. Singh, M. K. Reproducibility and Reliability of Computing Models in Segmentation and Volumetric Measurement of Brain. Ann Neurosci 30, 224–229 (2023).

21. Gong, T. et al. Neurometabolic timecourse of healthy aging. Neuroimage 264, (2022).

22. Hafkemeijer, A. et al. Associations between age and gray matter volume in anatomical brain networks in middle-aged to older adults. Aging Cell 13, 1068–1074 (2014).

23. Terribilli, D. et al. Age-related gray matter volume changes in the brain during non-elderly adulthood. Neurobiol Aging 32, 354–368 (2011).

24. Mahmoudi, N. et al. Microstructural and Metabolic Changes in Normal Aging Human Brain Studied with Combined Whole-Brain MR Spectroscopic Imaging and Quantitative MR Imaging. Clin Neuroradiol 33, 993–1005 (2023).

25. Maghsudi, H. et al. Age-related Brain Metabolic Changes up to Seventh Decade in Healthy Humans: Whole-brain Magnetic Resonance Spectroscopic Imaging Study. Clin Neuroradiol 30, 581–589 (2020).

26. Hui, S. C. N., et al. sLASER and PRESS perform similarly at revealing metabolite-age correlations at 3 T. Magn Reson Med 91, 431–442 (2024).

27. Jirsaraie, R. J. et al. Benchmarking the generalizability of brain age models: Challenges posed by scanner variance and prediction bias. Hum Brain Mapp 44, 1118–1128 (2023).

28. Archibald, J., Bouchard, A. E., Noeske, R., Shungu, D. C. & Mikkelsen, M. Test-retest reliability of multi-metabolite edited MRS at 3T using PRESS and sLASER. Preprint at 10.1101/2025.06.07.657685 (2025).

29. Deelchand, D. K. et al. Across-vendor standardization of semi-LASER for single-voxel MRS at 3T. NMR Biomed 34, (2021).

30. Tkáč, I., et al. Water and lipid suppression techniques for advanced 1H MRS and MRSI of the human brain: Experts’ consensus recommendations. NMR in Biomedicine vol. 34 Preprint at 10.1002/nbm.4459 (2021).

31. Oeltzschner, G. et al. Osprey: Open-source processing, reconstruction & estimation of magnetic resonance spectroscopy data. J Neurosci Methods 343, (2020).

32. Simpson, R., Devenyi, G. A., Jezzard, P., Hennessy, T. J. & Near, J. Advanced processing and simulation of MRS data using the FID appliance (FID-A)—An open source, MATLAB-based toolkit. Magn Reson Med 77, 23–33 (2017).

33. Landheer, K. & Juchem, C. Are Cramér-Rao lower bounds an accurate estimate for standard deviations in in vivo magnetic resonance spectroscopy? NMR Biomed 34, (2021).

34. Gasparovic, C., Chen, H. & Mullins, P. G. Errors in 1H-MRS estimates of brain metabolite concentrations caused by failing to take into account tissue-specific signal relaxation. NMR Biomed 31, (2018).

35. Mlynrik, V., Gruber, S. & Moser, E. Proton T1 and T2 relaxation times of human brain metabolites at 3 Tesla. NMR Biomed 14, 325–331 (2001).

36. Dhamala, E. et al. Validation of in vivo MRS measures of metabolite concentrations in the human brain. NMR Biomed 32, (2019).

37. Wyss, P. O. et al. In vivo estimation of transverse relaxation time constant (T2) of 17 human brain metabolites at 3T. Magn Reson Med 80, 452–461 (2018).

38. Stegmann, G., Jacobucci, R., Harring, J. R. & Grimm, K. J. Nonlinear Mixed-Effects Modeling Programs in R. Structural Equation Modeling vol. 25 160–165 Preprint at 10.1080/10705511.2017.1396187 (2018).

39. Steiger, J. H. Tests for comparing elements of a correlation matrix. Psychol Bull (1980).

40. Kruse, A. O. & Bustillo, J. R. Glutamatergic dysfunction in Schizophrenia. Translational Psychiatry vol. 12 Preprint at 10.1038/s41398-022-02253-w (2022).

41. Swanberg, K. M., Kurada, A. V., Prinsen, H. & Juchem, C. Multiple sclerosis diagnosis and phenotype identification by multivariate classification of in vivo frontal cortex metabolite profiles. Sci Rep 12, (2022).

42. Swanberg, K. M., Landheer, K., Pitt, D. & Juchem, C. Quantifying the Metabolic Signature of Multiple Sclerosis by in vivo Proton Magnetic Resonance Spectroscopy: Current Challenges and Future Outlook in the Translation From Proton Signal to Diagnostic Biomarker. Frontiers in Neurology vol. 10 Preprint at 10.3389/fneur.2019.01173 (2019).

43. Murray, M. E. et al. Early Alzheimer’s disease neuropathology detected by proton MR spectroscopy. Journal of Neuroscience 34, 16247–16255 (2014).

44. Kantarci K, W. S. P. S., et al. Risk of Dementia in MCI Combined Effect of Cerebrovascular Disease, Volumetric MRI, and 1 H MRS. Mayo Clinic www.neurology.org (2009).

45. Luque, E. M., et al. An Update on MR Spectroscopy in Cancer Management: Advances in Instrumentation, Acquisition, and Analysis. Radiology: Imaging Cancer vol. 6 Preprint at 10.1148/rycan.230101 (2024).

46. Kantarci, K. et al. Longitudinal 1H MRS changes in mild cognitive impairment and Alzheimer’s disease. Neurobiol Aging 28, 1330–1339 (2007).

47. Archibald, J. et al. Metabolite activity in the anterior cingulate cortex during a painful stimulus using functional MRS. Sci Rep 10, (2020).

48. Hasler G, van der V. J. T. T. M. N. S. J. D. WC. Reduced prefrontal glutamate/glutamine and gamma-aminobutyric acid levels in major depression determined using proton magnetic resonance spectroscopy. Arch Gen Psychiatry (2007).

49. Clarke, W. T., Stagg, C. J. & Jbabdi, S. FSL-MRS: An end-to-end spectroscopy analysis package. Magn Reson Med 85, 2950–2964 (2021).

50. Piechnik, S. K., Evans, J., Bary, L. H., Wise, R. G. & Jezzard, P. Functional changes in CSF volume estimated using measurement of water T2 relaxation. Magn Reson Med 61, 579–586 (2009).

51. Resnick, S. M., Pham, D. L., Kraut, M. A., Zonderman, A. B. & Davatzikos, C. Longitudinal Magnetic Resonance Imaging Studies of Older Adults: A Shrinking Brain. (2003).

52. Maddock, R. J. Statistical non-independence of brain metabolite concentrations whether normalized to creatine or water. Journal of Cerebral Blood Flow and Metabolism Preprint at 10.1177/0271678X241290018 (2024).

53. Prisciandaro, J. J., Zöllner, H. J., Murali-Manohar, S., Oeltzschner, G. & Edden, R. A. E. More than one-half of the variance in in vivo proton MR spectroscopy metabolite estimates is common to all metabolites. NMR Biomed 36, (2023).

54. Gasparovic, C., Chen, H. & Mullins, P. G. Errors in 1H-MRS estimates of brain metabolite concentrations caused by failing to take into account tissue-specific signal relaxation. NMR Biomed 31, (2018).

55. Harris, A. D., Puts, N. A. J. & Edden, R. A. E. Tissue correction for GABA-edited MRS: Considerations of voxel composition, tissue segmentation, and tissue relaxations. Journal of Magnetic Resonance Imaging 42, 1431–1440 (2015).

56. Mikkelsen, M., Singh, K. D., Brealy, J. A., Linden, D. E. J. & Evans, C. J. Quantification of γ-aminobutyric acid (GABA) in 1H MRS volumes composed heterogeneously of grey and white matter. NMR Biomed 29, 1644–1655 (2016).

57. Taki, Y. et al. A longitudinal study of gray matter volume decline with age and modifying factors. Neurobiol Aging 32, 907–915 (2011).

58. Hafkemeijer, A. et al. Associations between age and gray matter volume in anatomical brain networks in middle-aged to older adults. Aging Cell 13, 1068–1074 (2014).

59. Instrella, R. & Juchem, C. Uncertainty propagation in absolute metabolite quantification for in vivo MRS of the human brain. Magn Reson Med 91, 1284–1300 (2024).

