## supplementary material for "Benchmarking the Impact of Anatomical Segmentation on In Vivo Magnetic Resonance Spectroscopy"

**Table S1.** **Assessment of normality of the distributions of metabolite concentration estimates.** Shapiro-Wilk tests were performed across all datasets (ANTS, FSL, SPM), sessions (Ses1 and Ses2), and metabolites (tNAA N-acetylaspartate; tCr total creatine; tCho total choline; mI myo-inositol; Glu glutamate; Glx combined glutamate and glutamine.).

| Dataset | Session | Metabolite | Adjusted P-value |
| --- | --- | --- | --- |
| ANTS | 1 | Glu | >0.99999 |
| ANTS | 1 | Glx | > 0.99999 |
| ANTS | 1 | mI | > 0.99999 |
| ANTS | 1 | tCho | > 0.99999 |
| ANTS | 1 | tCr | 0.951 |
| ANTS | 1 | tNAA | > 0.99999 |
| ANTS | 2 | Glu | > 0.99999 |
| ANTS | 2 | Glx | 0.940 |
| ANTS | 2 | mI | > 0.99999 |
| ANTS | 2 | tCho | > 0.99999 |
| ANTS | 2 | tCr | > 0.99999 |
| ANTS | 2 | tNAA | > 0.99999 |
| FSL | 1 | Glu | > 0.99999 |
| FSL | 1 | Glx | > 0.99999 |
| FSL | 1 | mI | > 0.99999 |
| FSL | 1 | tCho | > 0.99999 |
| FSL | 1 | tCr | > 0.99999 |
| FSL | 1 | tNAA | > 0.99999 |
| FSL | 2 | Glu | > 0.99999 |
| FSL | 2 | Glx | > 0.99999 |
| FSL | 2 | mI | > 0.99999 |
| FSL | 2 | tCho | > 0.99999 |
| FSL | 2 | tCr | > 0.99999 |
| FSL | 2 | tNAA | > 0.99999 |
| SPM | 1 | Glu | > 0.99999 |
| SPM | 1 | Glx | > 0.99999 |
| SPM | 1 | mI | > 0.99999 |
| SPM | 1 | tCho | > 0.99999 |
| SPM | 1 | tCr | > 0.99999 |
| SPM | 1 | tNAA | > 0.99999 |
| SPM | 2 | Glu | > 0.99999 |
| SPM | 2 | Glx | 0.509 |
| SPM | 2 | mI | > 0.99999 |
| SPM | 2 | tCho | > 0.99999 |
| SPM | 2 | tCr | > 0.99999 |
| SPM | 2 | tNAA | > 0.99999 |

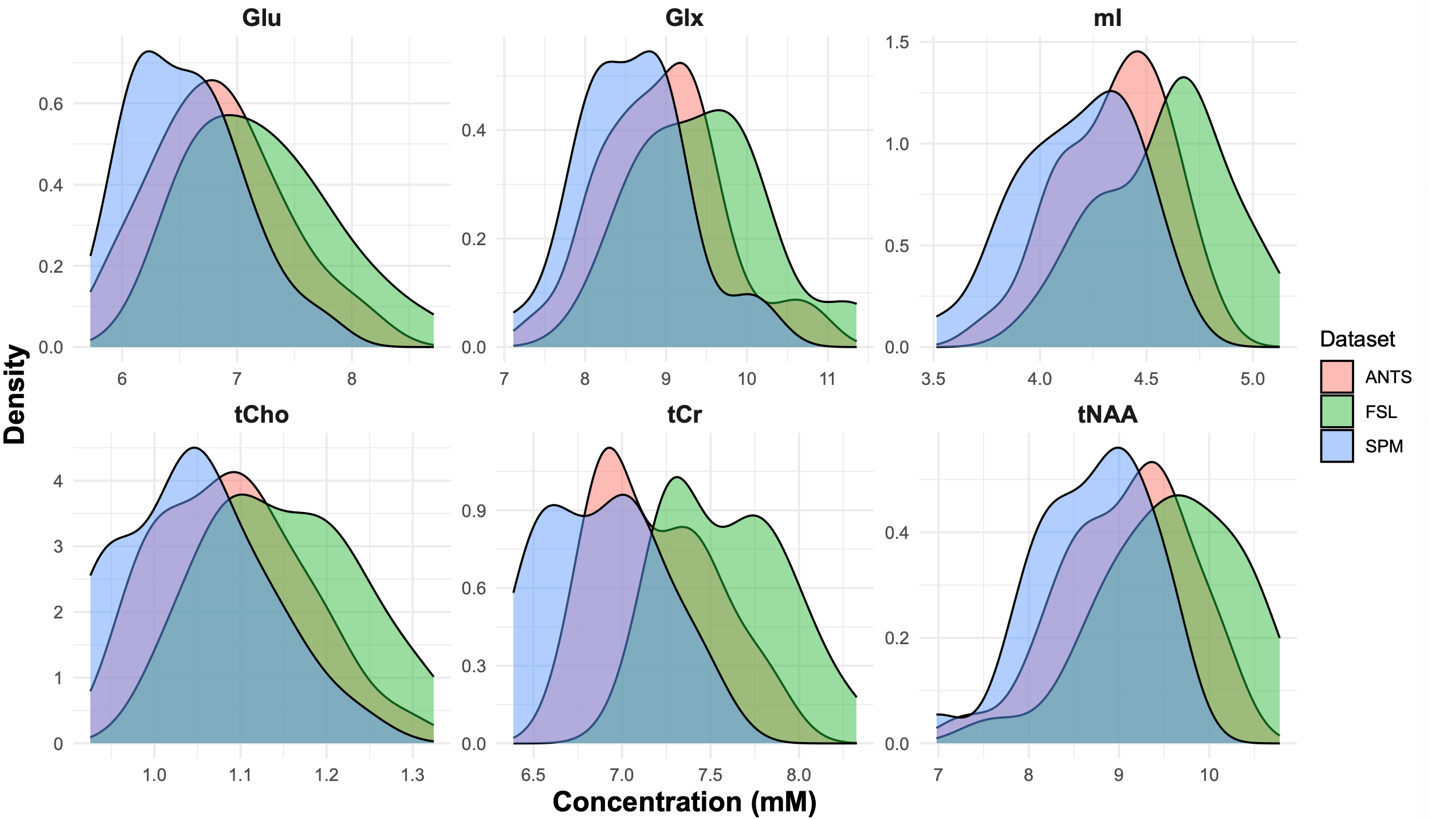

**Figure S1. Density plots of metabolite concentration values by dataset (segmentation method), faceted by metabolite.** Distributions are shown for each metabolite (e.g., tNAA, tCr, tCho, Glu, Glx, mI), with density curves overlaid for each segmentation method (ANTs, FSL, and SPM). These plots illustrate the variability and distributional characteristics across methods. Normality was statistically assessed using the Shapiro-Wilk test within each dataset, session, and metabolite, with Bonferroni-adjusted *p*-values reported.

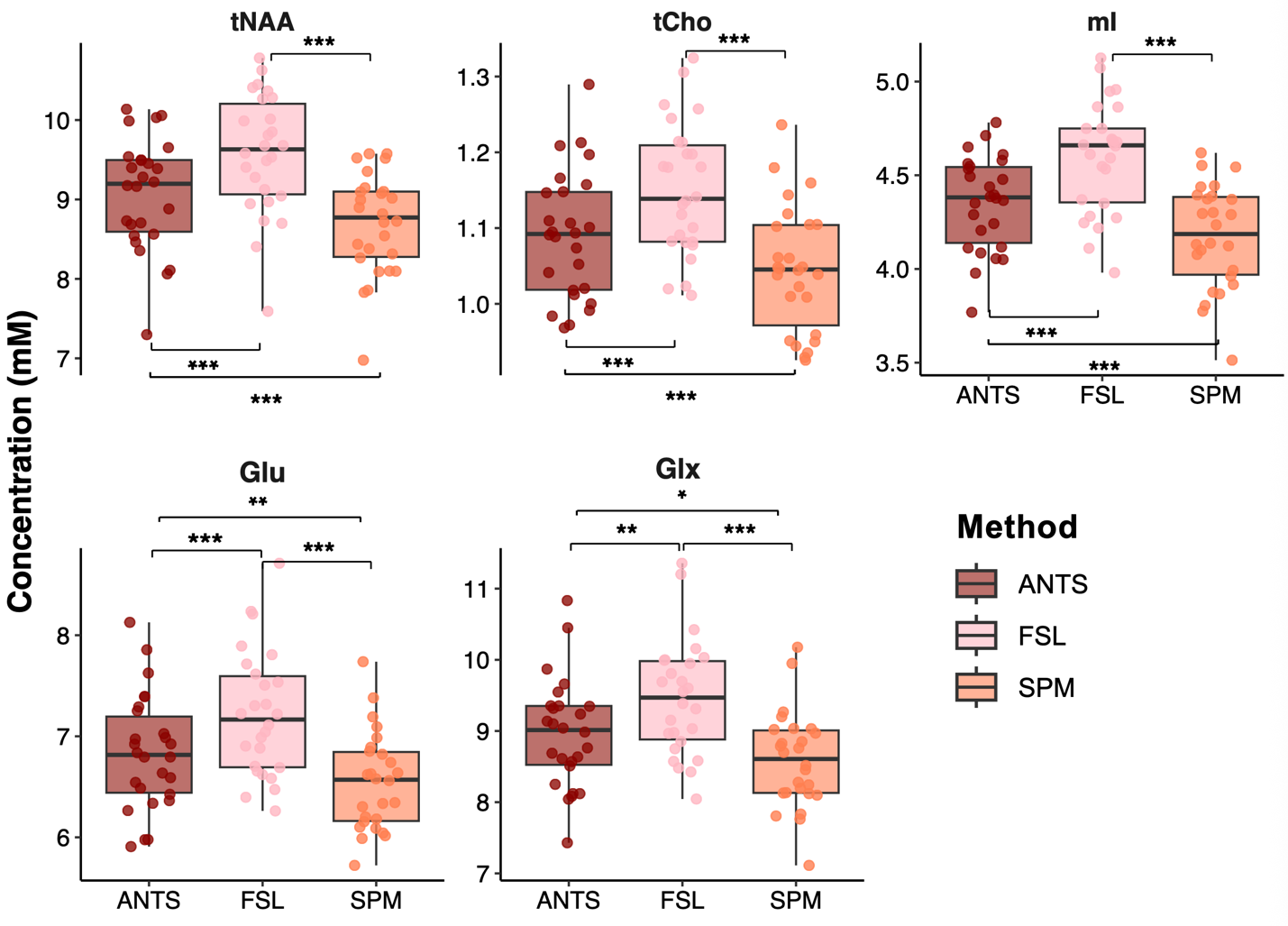

**Figure S2. Concentrations (mM) of metabolite levels across the three segmentation datasets (ANTs, FSL, and SPM).** Results are shown from a repeated-measures ANOVA with Tukey post hoc-corrected pairwise comparisons. Significant differences in metabolite concentration between datasets are indicated by asterisks (*p* < 0.05). tNAA, total *N*-acetylaspartate; tCho, total choline; mI, *myo*-inositol, Glu, glutamate; Glx, glutamate + glutamine. *p* < 0.05 = *, *p* < 0.01 = **, *p* < 0.001 = ***.

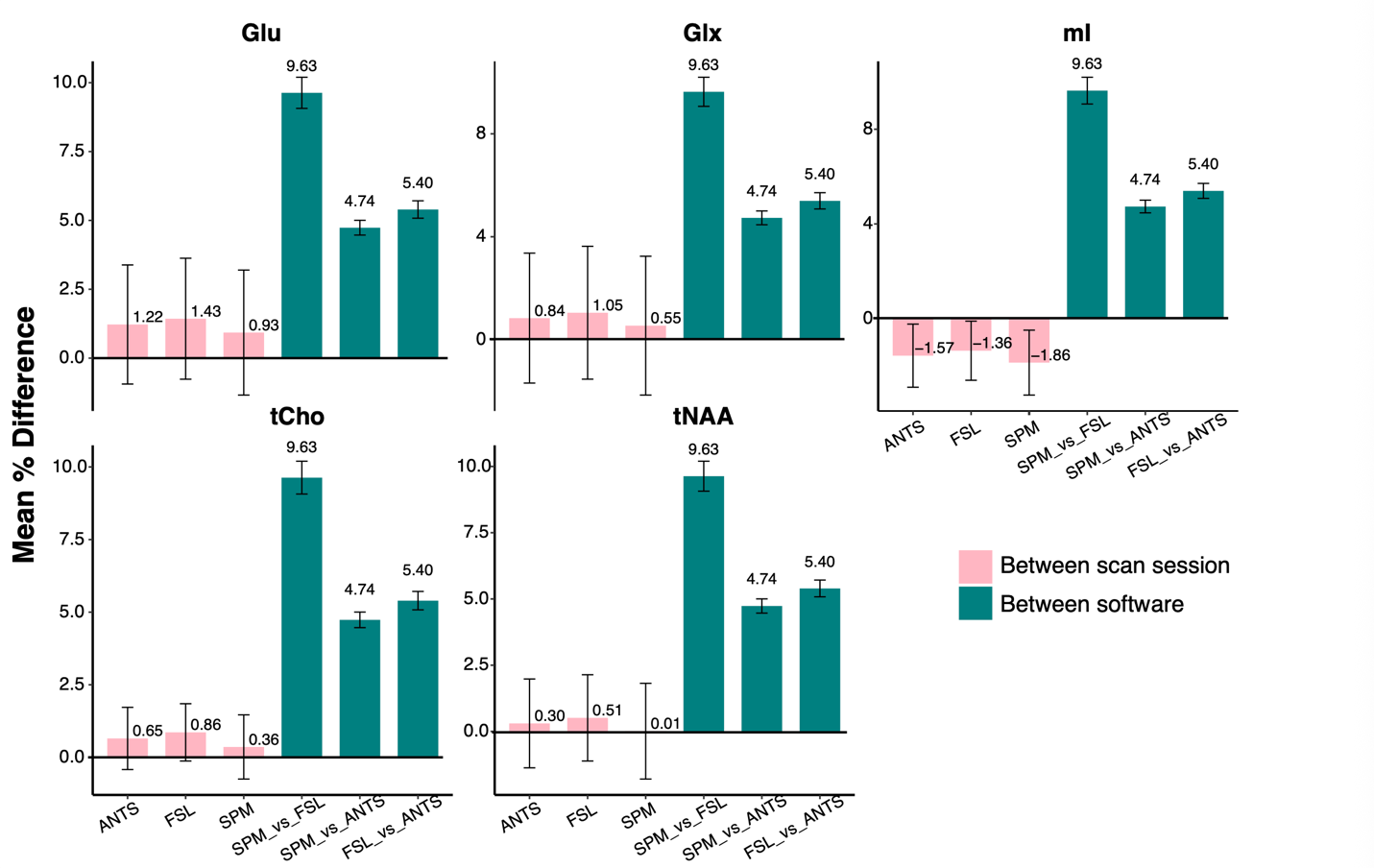

**Figure S3. Mean percentage differences for all metabolites across methods.** Bars represent the mean percentage difference between scan sessions (light pink) and between software packages (teal). Error bars denote the standard deviation. Glu, glutamate; Glx, glutamate + glutamine; mI, myo-inositol; tCho, total choline; tNAA, total *N*-acetylaspartate. *Note:* The similarity in percentage differences between software packages reflects the fact that the equation used to calculate molar concentrations impacts all metabolite estimates proportionally.

**Table S2.** **MRSinMRS checklist^1^**

| 1. Hardware |  |
| --- | --- |
| a. Field strength [T] | 3T |
| b. Manufacturer | GE |
| c. Model (software version if available) | Discovery MR750 (DV26.0_R01_1725.a) |
| d. RF coils: nuclei (transmit/ receive), number of channels, type, body part | ^1^H 32-channel phased-array head coil for receive; body coil for transmit |
| e. Additional hardware | n/a |
| 2. Acquisition |  |
| a. Pulse sequence | sLASER |
| b. Volume of Interest (VOI) locations | Medial parietal lobe |
| c. Nominal VOI size | 30 × 30 × 30 mm^3^ |
| d. Repetition Time (TR), Echo Time (TE) | TR = 2000 ms; TE = 35 ms |
| e. Total number of Excitations or acquisitions per spectrum | 64 |
| f. Additional sequence parameters | Spectral width: 5000 Hz  Spectral data points: 4096  GOIA‑WURST refocusing pulse; duration/bandwidth: 4.5 ms/10 kHz |
| g. Water Suppression Method | VAPOR; pulse duration/bandwidth: 30 ms/70 Hz |
| h. Shimming Method, reference peak, and thresholds for “acceptance of shim” chosen | Double-echo GRE |
| i. Triggering or motion correction method | n/a |
| 3. Data analysis methods and outputs |  |
| a. Analysis software | Osprey (v2.9.6); LCModel (embedded in Osprey) (v6.3-1N); R (v4.5.0) |
| b. Processing steps deviating from quoted reference or product | RF coil combination performed using generalized least squares (An et al., 2013, doi:[10.1002/jmri.23941](https://www.doi.org/10.1002/jmri.23941))  Basis set created for TE = 35 ms sLASER using FID-A (Simpson et al., 2017, doi:10.1016/j.neuroimage.2017.02.058), simulating 20 metabolites, LW = 2 Hz, spectral width = 5000 Hz, 64×64 grid. |
| c. Output measure  Processing steps deviating from quoted reference or product | Tissue- and relaxation-corrected “absolute” concentration estimates (mM) using internal tissue water as a concentration reference |
| d. Quantification references and assumptions, fitting model assumptions | Unsuppressed water used as a reference; model assumptions were as set by Osprey by default |
| 4. Data Quality |  |
| a. Reported variables  (SNR, Linewidth (with reference peaks)) | Creatine SNR; H_2_O FWHM; Fit Quality Index |
| b. Data exclusion criteria | Visual inspection of spectra |
| c. Quality measures of postprocessing Model fitting (e.g., CRLB, goodness of fit, SD of residual) | Fit error calculated as the sum of squares of residuals normalized to the square of the standard deviation of the noise signal between –2 and 0 ppm and multiplied by the number of points of the residuals |
| d. Sample spectrum | Provided in the main text of the article Figure 1 |

### **Table S3. Relaxation constants used in quantification calculations^2–4^.** Values are averaged over grey and white matter and metabolite moieties.

**(A) Metabolite *T*₁ and *T*₂ Values (ms)**

| Metabolite | *T*₁ (ms) | Reference | *T*₂ (ms) | Reference |
| --- | --- | --- | --- | --- |
| tNAA | 1410 | Mlynárik et al. (2001) | 282.25 | Wyss et al. (2018) |
| tCr | 1350 | Mlynárik et al. (2001) | 146.75 | Wyss et al. (2018) |
| tCho | 1190 | Mlynárik et al. (2001) | 241.71 | Wyss et al. (2018) |
| mI | 1090 | Mlynárik et al. (2001) | 202.5 | Wyss et al. (2018) |
| Glu | 1220 | Mlynárik et al. (2001) | 129.5 | Wyss et al. (2018) |
| Glx | 1080 | Mlynárik et al. (2001) | 129.5 | Wyss et al. (2018) |

#### **B) Water Relaxation Attenuation Factors**

| Tissue Type | *T*₁ (ms) | T₂ (ms) | *R*_H₂O,x_^*^ | Source |
| --- | --- | --- | --- | --- |
| Gray matter | 1331 | 110 | 0.565 | Dhamala et al. (2019) |
| White matter | 832 | 79.2 | 0.584 | Dhamala et al. (2019) |
| CSF | 3817 | 503 | 0.380 | Dhamala et al. (2019) |
| ^*^ Computed assuming TE/TR = 35/2000 ms | | | | |

**Table S4-** **Tukey-adjusted pairwise comparisons of metabolite levels across segmentation methods.** Significant main effects of segmentation method were observed for all metabolites: tNAA, *F*(2, 60) = 46.07, *p* < .0001; mI, *F*(2, 60) = 68.10, *p* < .0001; Glx, *F*(2, 60) = 21.79, *p* < .0001; Glu, *F*(2, 60) = 27.44, *p* < .0001; tCho, *F*(2, 60) = 88.63, *p* < .0001.There were no significant interactions between segmentation method and session for any metabolite (*all p* > 0.92), justifying the use of pairwise comparisons averaged across sessions

| Metabolite | Contrast | Mean diff | Std.  Error | df | t | p |
| --- | --- | --- | --- | --- | --- | --- |
| tNAA | ANTS – FSL | -0.51 | 0.09 | 60 | -5.47 | < .0001 |
|  | ANTS – SPM | 0.38 | 0.09 | 60 | 4.10 | 0.0004 |
|  | FSL – SPM | 0.88 | 0.09 | 60 | 9.57 | < .0001 |
| mI | ANTS – FSL | -0.24 | 0.04 | 60 | -6.70 | < .0001 |
|  | ANTS – SPM | 0.18 | 0.04 | 60 | 4.92 | < .0001 |
|  | FSL – SPM | 0.42 | 0.04 | 60 | 11.63 | < .0001 |
| Glx | ANTS – FSL | -0.50 | 0.13 | 60 | -3.75 | 0.0012 |
|  | ANTS – SPM | 0.38 | 0.13 | 60 | 2.84 | 0.0169 |
|  | FSL – SPM | 0.87 | 0.13 | 60 | 6.58 | < .0001 |
| Glu | ANTS – FSL | -0.38 | 0.09 | 60 | -4.20 | 0.0003 |
|  | ANTS – SPM | 0.29 | 0.09 | 60 | 3.18 | 0.0064 |
|  | FSL – SPM | 0.67 | 0.09 | 60 | 7.39 | < .0001 |
| tCho | ANTS – FSL | -0.06 | 0.01 | 60 | -7.60 | < .0001 |
|  | ANTS – SPM | 0.05 | 0.01 | 60 | 5.67 | < .0001 |
|  | FSL – SPM | 0.11 | 0.01 | 60 | 13.27 | < .0001 |

**References**

1.	Lin, A. *et al.* Minimum Reporting Standards for in vivo Magnetic Resonance Spectroscopy (MRSinMRS): Experts’ consensus recommendations. *NMR Biomed* 34, (2021).

2.	Mlynrik, V., Gruber, S. & Moser, E. Proton T1 and T2 relaxation times of human brain metabolites at 3 Tesla. *NMR Biomed* 14, 325–331 (2001).

3.	Wyss, P. O. *et al.* In vivo estimation of transverse relaxation time constant (T2) of 17 human brain metabolites at 3T. *Magn Reson Med* 80, 452–461 (2018).

4.	Dhamala, E. *et al.* Validation of in vivo MRS measures of metabolite concentrations in the human brain. *NMR Biomed* 32, (2019).

1. * Jessica Archibald and Kay Chioma Igwe contributed equally to this work
